# Association Between Vaginal Community States and Preeclampsia Status in Pregnant Individuals

**DOI:** 10.1101/2025.09.18.677097

**Authors:** Grace Ekalle, Sayumi York, Madeleine Gerard, Jennifer Kerr

## Abstract

Preeclampsia (PE) is a severe pregnancy complication affecting 3–8% of pregnancies. Recent evidence suggests the vaginal microbiome may influence PE risk. To investigate this, we reanalyzed publicly available 16S rRNA sequencing data (PRJNA798597) from vaginal samples of pregnant women with (n=10) and without PE (n=10) and assigned vaginal community state types which were previously uncharacterized. Overall microbial diversity did not differ significantly between groups; however, subtype differences in *Lactobacillus* spp. were observed. D-lactic acid-producing vaginal community state types were less common in PE, suggesting potential value for microbiome-based risk assessment.

## Description

Hypertensive disorders of pregnancy, including preeclampsia, are among the leading causes of maternal and fetal morbidity and mortality worldwide (Deady et al., 2024). Each year, over half a million women die from pregnancy-related causes, with 99% of these deaths occurring in low- and middle-income countries (Duley, 2009). Despite global advancements in healthcare, the underlying mechanisms driving these disorders remain unclear. Preeclampsia, which commonly occurs in first pregnancies, presents with variable symptoms but is typically defined by hypertension and proteinuria caused by stress-related placental factors (Duley, 2009).

Emerging evidence highlights the microbiome as a potential contributor to preeclampsia development. Dysbiosis, or imbalance within microbial communities, has been associated with adverse health outcomes in multiple systems, including cardiovascular and reproductive health (Kell & Kenny, 2016). While the gut microbiome has been extensively studied, the vaginal microbiome also plays an important role, particularly through vaginal community state types (CSTs) that influence pregnancy outcomes (Kell & Kenny, 2016).

CSTs classify vaginal microbial communities commonly observed in reproductive-age women (Ravel et al., 2011). Five CSTs have been described and recognized, four dominated by different *Lactobacillus* species and one characterized by a more balanced mix of facultative and obligate anaerobes (France et al., 2020). CST I, dominated by *L. crispatus*, is usually linked to vaginal health and resistance to infections. CST III, on the other hand, is dominated by *L. iners* and often shows higher levels of inflammatory cytokines such as TNF-α and IL-18 (Chee et al., 2020). This suggests that *L. iners*-dominated communities may be more vulnerable to shifts that could be associated with pregnancy complications.

CSTs can further be stratified into D-lactic acid–producing and non-D-lactic acid-producing groups (Plummer et al., 2021). Lactic acid is produced in the vagina by *Lactobacillus* species and exists in two forms, D- and L-lactic acid, depending on the strain of *Lactobacillus*. Species such as *L. crispatus* and *L. gasseri* produce both isomers, while *L. jensenii* produces only D-lactic acid and *L. iners* only L-lactic acid, which may partly explain the stronger protective role of *L. crispatus* compared to *L. iners*. D-lactic acid in particular is hypothesized to provide greater protection against upper genital tract infections than L-lactic acid, making it a possible factor in vaginal health and pregnancy outcomes.

A previous study by Geldenhuys et al.(2022) examined the vaginal microbiomes of 21 pregnant women in South Africa. Within the rarefied subset of 10 vaginal samples processed in QIIME 2 (pre-eclampsia, n=5; normotensive, n=5), pre-eclampsia was associated with higher alpha diversity and a reduced relative abundance of *Lactobacillus* spp. Women with pre-eclampsia exhibited significantly higher vaginal microbiome diversity and reduced *Lactobacillus* spp., although *Lactobacillus iners* remained predominant across both groups. Although differences in relative abundance were observed across multiple taxonomic ranks, the overall community structure did not differ significantly between groups. Species-level taxonomic assignments were not available at the time of publication, and per-sample community profiles were not reported, which limits the granularity and generalizability of these findings.

Our study builds on Geldenhuys et al. (2022) by applying VALENCIA, a vaginal microbiome specific tool, to all sampled vaginal microbiome samples (n=10 from both preeclamptic and normotensive individuals) to classify vaginal microbial communities at the species level and organize them into CSTs to assess potential links between vaginal microbiome subtypes and pre-eclampsia risk (Figure 1A). During this process we also classified sequences into ASVs, which are more precise than the OTU’s originally used, and can be replicated between studies (Callahan et al., 2017). Our findings establish a possible connection between preeclampsia risk and non-lactic producing acid CSTs for future studies to investigate.

**Figure 1.**
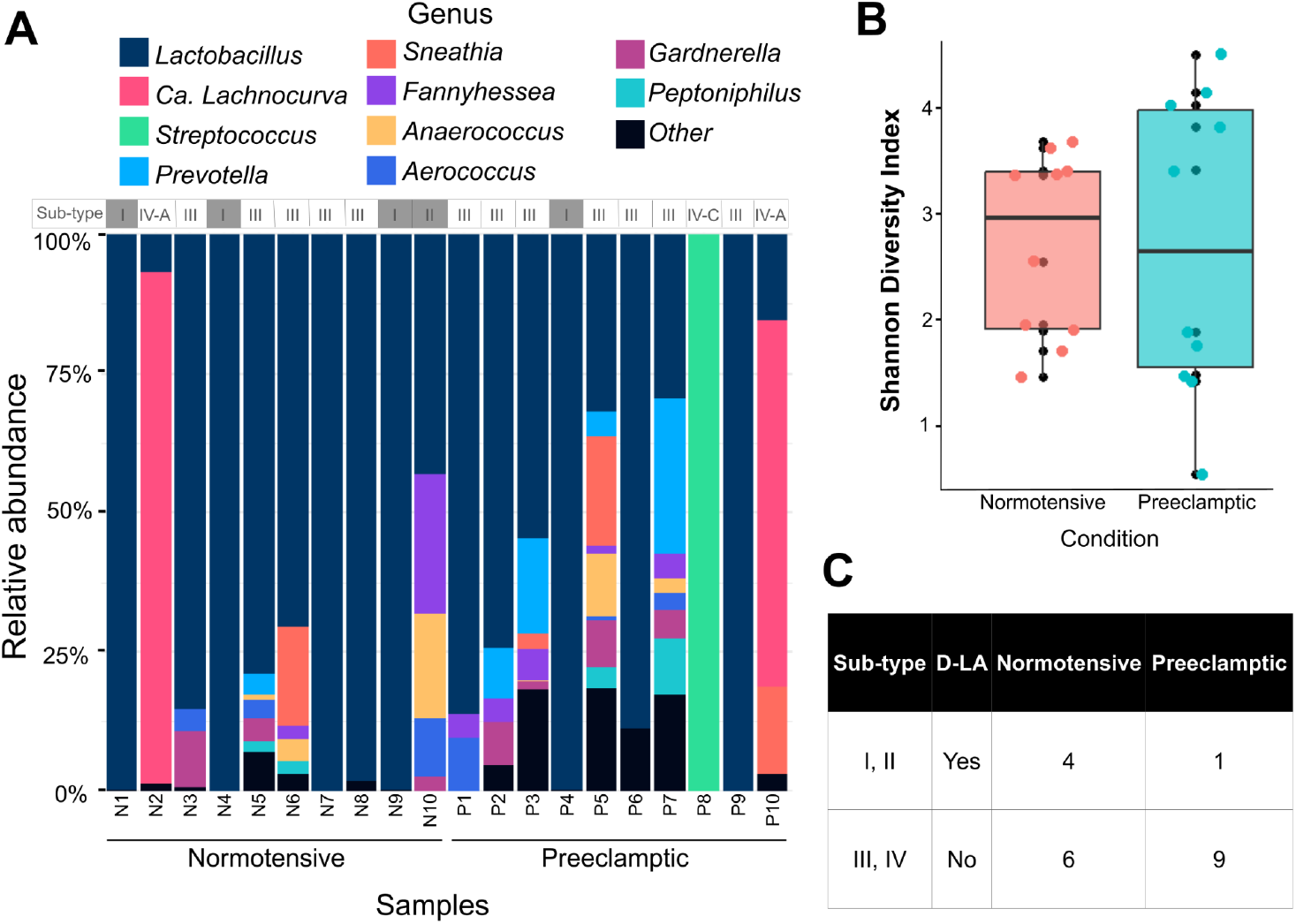
Vaginal microbial composition, diversity, and characterization of normotensive (n=10) and preeclamptic (n=10) individuals. **A**. Differential abundance of microbial taxa between normotensive and preeclamptic samples normalized by proportion. Individual community state type (CST) sub-classification shown. **B**. Shannon alpha-diversity richness measure comparing normotensive and preeclamptic samples. Line in box plot indicates median value. **C**. CST classification of normotensive and preeclamptic samples grouped into D-lactic acid–producing and non–D-lactic acid–producing CSTs, identified using VALENCIA.

342 ASVs were recovered across all samples. An average number of 21.7±16.4 ASVs (Range: 3-45) and 38.5±44.4 ASVs (Range: 1-132) were found in normotensive and preeclamptic samples respectively. We did not find a significant difference in the number of distinct ASVs between groups (p = 0.88). The difference between the average Shannon diversity of vaginal microbiomes in normotensive (2.5 ± 1.0) and preeclamptic groups (2.5 ± 1.6) was not found to be significantly different (p=0.82) (Figure 1B).

Samples N1, N4, N7, N8, N9, P4, P6, and P9 were dominated by the *Lactobacillus* genus, with no notable variation in microbial composition. These samples primarily clustered into CST I and CST III (Figure 1A). Samples N2 and P10 were heavily dominated by *Candidatus Lachnocurva vaginae*, with additional contributions from *Lactobacillus* spp. (N2 and P10) and *Sneathia* spp. (P10). Both samples were classified as CST IV-A. Sample P8 exhibited a strong predominance of the *Streptococcus* genus and was assigned to CST IV-C. Samples N3, N5, N6, N10, P1, P2, P3, P5, and P7 demonstrated greater microbial diversity overall, though *Lactobacillus* spp. remained the dominant taxa.

CST subtype differences in *Lactobacillus* spp. were observed across groups. Notably, vaginal community subtype analysis revealed that samples classified as D-lactic acid–producing CSTs were 80% less likely to belong to the pre-eclampsia group, although this trend did not reach statistical significance (95% CI: 0.01–1.86) (Figure 1C).

This study aimed to classify vaginal microbial communities at the species level using VALENCIA and to assess whether CST differences, particularly D-lactic acid-producing versus non-D-lactic acid-producing groups, were linked to preeclampsia risk. While observed and alpha diversity metrics were not significantly different between normotensive and preeclamptic groups, species-level CST classification revealed possible differences between normotensive and preeclamptic groups.

Overall, our study supports previous findings that suggest community composition is a more reliable predictor of pregnancy outcomes than overall diversity (DiGiulio et al., 2015). Vaginal microbiome diversity differs by ethnicity (Hyman et al., 2014). Our subjects, primarily of African ancestry, had similar Shannon diversity indices as subjects in other studies of the same ethnicity (Hyman et al., 2014). While we included all ASVs in our analysis, other studies may choose to exclude rare taxa (Aagaard et al., 2012), making direct comparisons difficult.

D-lactic acid-producing CST’s I and III were more frequent among normotensive subjects than preeclamptic subjects, suggesting a possible protective effect against preeclampsia. We classified 3 samples, 1 normotensive and 2 preeclamptic samples as CST IV. CST IV subtypes contained BV-associated taxa such as *Candidatus Lachnocurva vaginae* (BVAB1) and *Streptococcus*. Previous studies have linked BV-associated bacteria such as BVAB1 to higher risks of adverse outcomes, including spontaneous preterm birth, especially in women with prior preterm Birth (Nelson et al., 2014, 2015). The presence of BVAB1 may promote inflammation, weaken epithelial integrity, contribute to unrecognized infection, and disrupt immune defenses, all of which could contribute to preeclampsia (Gerede et al., 2024; Haggerty et al. 2008). *Prevotella* species, another CST IV member, are known to trigger inflammation through cytokine production and have been associated with premature rupture of membranes and preterm birth (Gerede et al., 2024). Their ability to produce virulence factors may compromise cervical tissue, creating vulnerability to hypertensive conditions and early labor.

Another genus frequently detected in our dataset, particularly among preeclamptic participants, is *Fannyhessea* (formerly *Atopobium*), whose presence typically marks a shift away from *Lactobacillus* dominance to a more varied vaginal ecosystem. A study by Odogwu et al. in 2021 reported that individuals who delivered preterm more often had vaginal communities dominated by *Fannyhessea (Atopobium) vaginae* than those delivering at term, and a higher midtrimester relative abundance of *F. (A*.*) vaginae* strongly predicted preterm birth, underscoring its potential as an early microbial risk marker (Odogwu et al., 2021).

More than half of normotensive samples were also associated with non-D-lactic acid-producing CSTs. Larger studies are needed to confirm the specificity of this association. It is also important to note that the small sample size in this study restricted statistical power and generalizability. Additionally, nine of the twenty samples contained fewer than ten ASVs, and one contained only a single ASV, raising concerns about sequencing depth. Samples that are expected to contain high amounts of host DNA, including vaginal samples can also be expected to have lower microbial sequencing yield (Pereira-Marques et al., 2019). Removal of host DNA via target enrichment sequencing technology has previously been used to successfully increase microbial sequencing depth from vaginal samples (Marquet et al. 2022).

Preeclampsia remains a leading contributor to medically indicated preterm birth due to the risks it poses to both mother and fetus. Reduced *Lactobacillus* levels with increased abundance of *Prevotella, Gardnerella*, and *Sneathia*, may amplify inflammation and weaken cervical integrity (Fettweis et al., 2019). This suggests that changes in microbiome composition may represent a biological pathway linking preeclampsia to preterm labor.

By classifying samples into CSTs, we found that preeclampsia cases were nearly always associated with non-D-lactic acid-producing CSTs. Since D-lactic acid plays a key role in protecting the vaginal barrier and regulating immune responses (Gerede et al., 2024), this may point to a functional link between microbial metabolism and hypertensive disorders of pregnancy. This work establishes a foundation for exploring CSTs as potential biomarkers for preeclampsia risk, facilitates future comparisons between studies through generating ASVs, and adds to the current body of knowledge of the vaginal microbiome and its associations with hypertensive disorders of pregnancy.

## Acknowledgements

I would like to express my gratitude to Gugulethu Sakana and Gauri Paul for their support throughout the research process and to C-MOOR for guidance with coding, troubleshooting, and organization of this microPublication. Thank you to Johanna Holm for her assistance with analyzing the vaginal state type grouping. This research was supported through grant UE5HG013799-01 and the Maryland E-Nnovation Initiative Fund.

## Methods

Raw data was downloaded from NCBI SRA (BioProject PRJNA798597) and imported into Galaxy for processing using a published dada2 workflow on Galaxy Training with modifications for single-end reads (Batut, 2025; Batut et al., 2018; Blankenberg et al., 2014; Hiltemann et al., 2023). In brief, sequence quality was assessed with dada2’s *plotQualityProfile* function and fastp (Callahan et al., 2016; Chen et al., 2018). Reads were trimmed to 450bp and reads with more than 2 expected errors were removed. The *dada* function of dada2 was then used to infer sequence variants that were subsequently evaluated with the dada2 *makeSequenceTable* function and *dada2*:*removeBimeraDenovo* to remove chimeras. Taxonomy assignment performed using *dada2:assignTaxonomy* and the learnErrors function rates with the gtdb_2018_11 database (Parks et al., 2018, 2020, 2022; Rinke et al., 2021). The SpeciateIT algorithm was used to determine additional vaginal microbiome genera and species from the final ASVs with the June 4, 2024 Speciate DB V3V4 model (Holm et al., 2024). Finally, results were imported and analyzed in the R package *phyloseq* using the and visualized with *ggplot2* (McMurdie & Holmes, 2013; Wickham, 2011).

We calculated the Shannon Diversity Index as a measure of sample alpha diversity using phyloseq’s *estimate_richness function*. A Kruskal-Wallis test was then used to compare the average difference in the number of observed ASVs and average Shannon Index between the normal and preeclamptic groups. Samples were classified into vaginal microbiome community state types (CSTs) using VALENCIA (France et al., 2020), a nearest-centroid based algorithm using the August 19, 2024 reference centroids and Python 3.0+. CSTs were grouped into D-Lactic acid producing state types (I, II) and non D-Lactic acid (III, IV) types. The oddsratio function from the epitools R packages was used to determine the odd ratio with 95% confidence interval (Aragon, 2004).

